# Complement receptor 1 gene (*CR1*) intragenic duplication and risk of Alzheimer’s disease

**DOI:** 10.1101/284711

**Authors:** Ezgi Kucukkilic, Keeley Brookes, Imelda Barber, Tamar Guetta-Baranes, Kevin Morgan, Edward J Hollox

## Abstract

Single nucleotide variants (SNVs) within and surrounding the complement receptor 1 (*CR1*) gene show some of the strongest genome-wide association signals with late-onset Alzheimer’s disease. Some studies have suggested that this association signal is due to a duplication allele (*CR1*-B) of a low copy repeat (LCR) within the *CR1* gene, which increases the number of complement C3b/C4b-binding sites in the mature receptor. In this study, we develop a triplex paralogue ratio test (PRT) assay for *CR1* LCR copy number allowing large numbers of samples to be typed with a limited amount of DNA. We also develop a *CR1*-B allele-specific PCR based on the junction generated by an historical non-allelic homologous recombination event between *CR1* LCRs. We use these methods to genotype *CR1* and measure *CR1-*B allele frequency in both late-onset and early-onset cases and unaffected controls from the United Kingdom. Our data support an association of late-onset Alzheimer’s disease with the *CR1*-B allele, and confirm that this allele occurs most frequently on the risk haplotype defined by SNV alleles. Furthermore, regression models incorporating *CR1*-B genotype provide a bitter fit to our data compared to incorporating the SNP-defined risk haplotype, supporting the *CR1*-B allele as the variant underlying the increased risk of late-onset Alzheimer’s disease.

## Introduction

Alzheimer’s disease (AD) is a common neurodegenerative disease with an increasing disease burden in an aging population (Ballard et al. 2011). Familial early-onset AD can be caused by autosomal dominant variants in, for example, the amyloid precursor protein gene *APP*, the presenilin 1 gene *PSEN1*, and the presenilin 2 gene *PSEN1* (Campion et al. 1999). Sporadic early onset AD (EOAD), with an age-of-onset of 65 years or less, is defined as disease in the absence of these classical familial early-onset AD mutations. However, 99% of AD cases are late-onset AD (LOAD), which is a complex disease with multiple environmental and genetic contributions to its etiology. The most important genetic variant affecting LOAD risk is the *APOE*\*E4 allele, which is a haplotype formed by rs429358-C and rs7412-C, generating a protein called APOE4 carrying arginine residues at position 130 and position 176 (Corder et al. 1998; Corder et al. 1993). This variant is associated with a 2-3-fold increase in LOAD risk for carriers, and a 15-fold increase in risk for individuals homozygous for this variant (Farrer et al. 1997). Genome-wide association studies (GWAS) on large cohorts have robustly identified 23 other variants with smaller effect sizes (Lambert et al. 2013; Lambert et al. 2009). Many of the genetic associations are with variants that lie within or near genes involved in the immune response. This has highlighted the importance of the immune response to amyloid plaque formation in Alzheimer’s disease, possibly mediated by microglial cells (Efthymiou and Goate 2017; Naj and Schellenberg 2017; Villegas-Llerena et al. 2016).

One of the genes implicated in LOAD risk by GWAS is the complement C3b/C4b receptor 1 gene *CR1*. The receptor encoded by this gene is expressed on the surface of leukocytes and erythrocytes, and binds the C3b fragment of complement C3, and the C4b fragment of complement C4, as well as complement C1q. These interactions are important in the clearance of antibody-antigen immune complexes from the blood circulation, and in the phagocytosis of complement-tagged pathogens. Complement receptor 1 is also involved in the inflammatory response to injured tissue (Holers 2014).

Alleles at several SNVs both proximal, distal and within the *CR1* gene have been identified as associated with LOAD, and these alleles are on a single risk haplotype that spans the *CR1* gene (Corneveaux et al. 2010; Lambert et al. 2009; Luo et al. 2014). Identifying the variant within this haplotype that is functionally responsible for the genetic association is challenging, yet correct identification will allow a functional genetic approach to determine the consequences of the variation in CR1 function, and therefore how that variation contributes to LOAD risk. As previously observed, there are at least 60 missense variants within the risk haplotype (Corneveaux et al. 2010). and it has been suggested that a rare missense variant rs4844609 might be responsible for the observed association (Keenan et al. 2012) but this observation has not been supported (Van Cauwenberghe et al. 2013).

*CR1* is known to contain an intragenic copy number variant (CNV) that alters the number of exons while maintaining the reading frame of the protein. The copy number variant is due to variable numbers of a tandemly-arranged 18kb repeat unit called a low copy repeat (LCR) (Crehan et al. 2012; Vik and Wong 1993; Wong et al. 1989; Wong et al. 1983). Each LCR contains eight exons, which together encode a C3b/C4b binding domain such that high copy numbers of the LCR result in a longer *CR1* molecule with more C3b/C4b binding domains (Figure 1). The *CR1* CNV has four alleles: *CR1*-A with two LCRs, *CR1*-B with three LCRs, *CR1*-C with one LCR and *CR1*-D with four LCRs. Each allele can be therefore represented by the copy number of the LCR domains. Alleles *CR1*-A, *CR1*-B and *CR1*-C are also known as *CR1*-F, *CR1*-S and *CR1*-F’ respectively in the literature, based on their mobilities in protein electrophoresis. Previous studies have shown that *CR1*-A is the most frequent allele in individuals of European origin with a frequency of 0.87. *CR1*-B is the next most frequent, at a frequency of 0.11, with *CR1*-C and *CR1*-D at frequencies of 0.02 and <0.01 respectively (Moulds et al. 1996).

**Figure 1.**
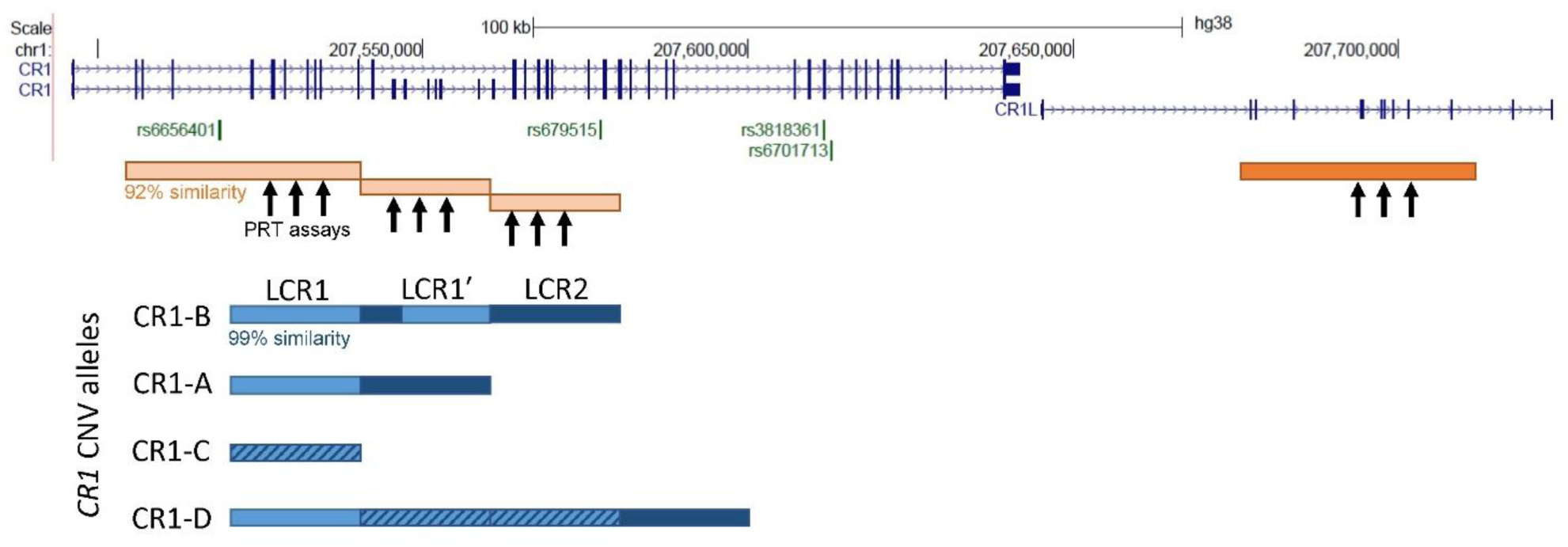
Structure of the human *CR1* and *CR1L* region showing the different CNV alleles. The *CR1* and *CR1L* genes are shown, with SNVs that have been reported as associated with late-onset Alzheimer’s disease in GWAS shown in green. The duplicated structure of the region is shown, with pale/dark orange boxes showing repeated regions that are ∼92% similar, and pale/dark blue boxes showing regions that are ∼99% similar. The genome assembly shows a *CR1*-B allele, comprised of LCR1, LCR2 and LCR1’ which is a fusion of LCR2 and LCR1. The structure of the alternative alleles (*CR1*-A, *CR1*-C and *CR1*-D) are shown below, with hatching indicating unclear origin of the LCR. The location of the amplification products for the three PRT assays are indicated, with the test amplicons generated from the pale orange repeats, and the reference amplicons generated from the dark orange repeat. Based in part on the UCSC Genome browser hg38 assembly.

Although the *CR1* LCR CNV affects protein sequence quite dramatically, it is not directly assayed by current GWAS. Nevertheless, the association signal observed with SNV haplotypes might be due to a *CR1* LCR CNV allele if that allele was on that particular SNV haplotype and therefore in linkage disequilibrium with the SNV alleles that show association with LOAD. A previous study tested the association of the *CR1*-B allele with LOAD on a cohort of Flemish Belgian patients (n=1039) and controls (n=844), and showed that *CR1*-B carriers showed an increased risk of LOAD (OR 1.32, 95% CI 1.03-1.69, p=0.028). They replicated their result on a sample of French patients (n=1393) and controls (n=610) (OR=1.33, 1.02-1.74, p=0.039), and showed that this represented the same association signal as that for the SNVs rs4844610 and rs1408077 (Brouwers et al. 2012).

Our study aimed firstly to develop a robust assay based on the paralogue ratio test and junction-fragment PCR for *CR1* LCR CNV, in particular for the intragenic duplication allele *CR1*-B, to facilitate further studies. Secondly, we aimed to use our methods to replicate the previous association of *CR1* and LOAD in a larger cohort. We also investigate the association of *CR1*-B in a cohort of EOAD to test for a stronger risk effect as a result of a more pronounced phenotype.

## Methods

### Samples

Lymphoblastoid cell lines derived from the 1000 Genomes project samples (Coriell Cell repositories) were grown using the standard conditions recommended by the supplier, and DNA isolated using a standard phenol-chloroform-based approach. UK samples were purchased as part of Human Random Controls plate 1 (HRC-1) from Public Health England.

Human DNA samples were obtained from the Alzheimer’s Research Trust (ART) Collaboration (University of Nottingham; University of Manchester; University of Southampton; University of Bristol; Queen’s University, Belfast; the Oxford Project to Investigate Memory and Ageing [OPTIMA], Oxford University); All case samples were diagnosed as either definite (post mortem confirmed) or probable Alzheimer’s disease (AD) according to National Institute of Neurological and Communicative Disorders and Stroke and the Alzheimer’s Disease and Related Disorders Association (NINCDS-ADRDA), and the Consortium to Establish a Registry for Alzheimer’s Disease (CERAD) guidelines. All samples used in this study were received with informed consent and were approved by the local Ethics Committee.

The 449 sporadic EOAD samples had an age of disease onset (AAO) ≤ 65 years of age, the 1436 LOAD samples had an AAO > 65 years of age and the 1359 controls had an age at death (AAD) > 65 years of age. Where AAO was not documented, it was derived assuming 8 years disease duration from age at death (Brookmeyer et al. 2002), or age at sampling (AAS) was used with the understanding it would approximate to disease onset (Tables 1 and 3).

**Table 1.**
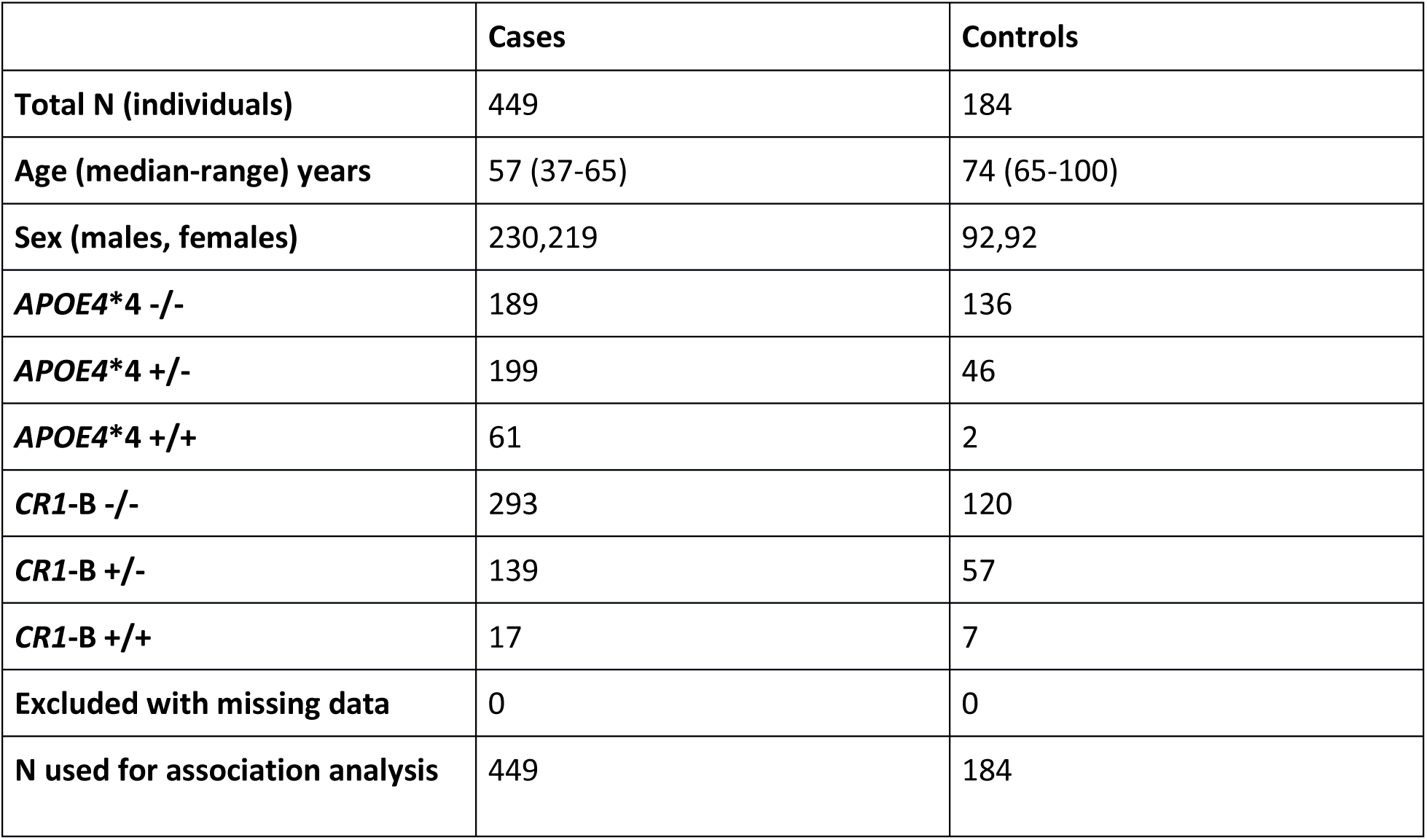
Characteristics of EOAD cohort and controls.

**Table 2.**
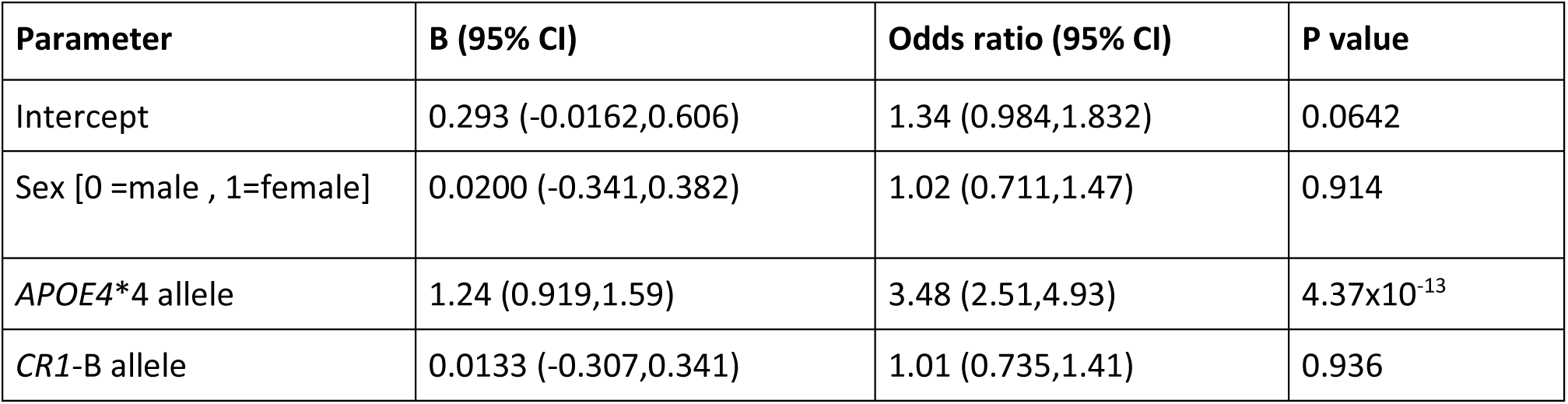
Association of *CR1*-B allele with EOAD.

| Parameter | B (95% CI) | Odds ratio (95% CI) | P value |
| --- | --- | --- | --- |
| Intercept | 0.293 (-0.0162,0.606) | 1.34 (0.984,1.832) | 0.0642 |
| Sex [0 =male , 1=female] | 0.0200 (-0.341,0.382) | 1.02 (0.711,1.47) | 0.914 |
| *APOE4* *4 allele | 1.24 (0.919,1.59) | 3.48 (2.51,4.93) | 4.37x10<sup>-13</sup> |
| *CR1*-B allele | 0.0133 (-0.307,0.341) | 1.01 (0.735,1.41) | 0.936 |

**Table 3.** Characteristics of LOAD cohort and controls.

|  | **Cases** | **Controls** |
| --- | --- | --- |
| **N (individuals)** | 1436 | 1175 |
| **Age (median-range) years** | 75.1 (43-98) | 73.2 (30-100) |
| **Sex (males, females)** | 569, 851 | 532, 639 |
| ***APOE4*\*4 -/-** | 605 | 863 |
| ***APOE4*\*4 +/-** | 670 | 286 |
| ***APOE4*\*4 +/+** | 161 | 26 |
| ***CR1*-B -/-** | 935 | 839 |
| ***CR1*-B +/-** | 422 | 284 |
| ***CR1*-B +/+** | 79 | 52 |
| **Excluded with missing age/sex data** | 256 | 170 |
| **N used for association analysis** | 1180 | 1005 |

DNA was extracted from blood or brain tissue using a standard phenol chloroform extraction method. DNA quality and quantity was assessed via gel electrophoresis and NanoDrop™ 3300 spectrometer respectively.

### CR1 copy number estimation from sequence read depth data

Sequence alignment files in .bam format and corresponding index files in .bai format were downloaded from the European Bioinformatics Institute (http://ftp.1000genomes.ebi.ac.uk/vol1/ftp/data_collections/1000_genomes_project/). Using samtools software (using the command `samtools view -c -F 4 input.bam target_region`), the number of mapped reads was counted across two intervals (GRCh37 chr1:207,697,239-207,751,921 test region, GRCh37 chr1:207,953,949-208,008,574 reference) for each of the samples analysed as part of the 1000 Genomes project (CEU individuals for Salt Lake City, Utah, Chinese individuals from Beijing, Japanese individuals for Tokyo and Yoruba individuals from Ibadan, Nigeria). A ratio of the reads from the copy number variable region:non-copy number variable region was taken as an estimate of *CR1* intragenic copy number. Data are available in dbvar accession nstd159 at ncbi.nlm.nih.gov/dbvar.

### CR1 paralogue ratio test

To determine *CR1* intragenic copy number on large numbers, we decided to design three specific assay using the paralogue ratio test (PRT), a form of competitive PCR that amplifies a test and reference locus using common primers, and uses the ratio of test and reference amplification products as a measure of copy number (Armour et al. 2007; Hollox 2017) (Supplementary Table 1). For each PCR, 5-10ng genomic DNA was amplified in a final volume of 10µl, containing 0.5 units *Taq* DNA polymerase (KAPA), 0.5µl of 10µM forward primer, 0.5µl of 10µM reverse primer and 1 µl of 10xPCR mix (10xPCR mix = 50mM TrisHCl pH8.8@25°C, 12.5mM ammonium sulphate, 1.4mM MgCl_2_, 125µg/ml BSA (Ambion), 7.5mM 2-mercaptoethanol and each dNTP (Promega) at a concentration of 200µM). Following PCR amplification in an Applied Biosystems Veriti thermal cycler at 95°C for 2 minutes, followed by between 24 and 27 cycles of 95°C for 20 seconds, 61-63°C for 20 seconds and 72°C for 20 seconds, and finally an elongation step at 72°C for 10 minutes. Cycle number and annealing temperatures for each assay are given in supplementary table 1.

Between 0.5-1µl of the final product from each assay was combined and added to HiDi Formamide (Applied Biosystems) containing 1% MapMarker ROX-labelled size standard. Following denaturation of the mixture at 96°C for 3 minutes and snap cooling on ice, the products were run on an Applied Biosystems 3130xl capillary sequencer following the manufacturer’s instructions, and areas of the test and reference peaks recorded using Genemapper software. For each assay, seven samples of known *CR1* LCR copy number were analysed with each experiment (Supplementary table 2). These samples were chosen from the HapMap phase I panel, with copy number inferred from previous array CGH data (Conrad et al. 2009) or from preliminary experiments with multiple PRTs. PRT values were normalised using the values from the seven positive controls to generate an estimated copy number value.

### Calling integer copy number from CR1 PRT data

Data from PRT1, PRT2 and PRT3 were concordant across the 1000 Genomes samples analysed so an average was taken to represent copy number. For each cohort, a Gaussian mixture model of four or five components was fitted to the data using the CNVtools software implemented in the statistical language R v.3.2.3, with each component representing a integer copy number class (Barnes et al. 2008). Samples were then assigned to each component with a posterior probability, and the component to which they were assigned reflected integer copy number call. A statistic Q, which is defined as the ratio of the separation of adjacent component means divided by the within-component standard deviation, averaged across all components of the mixture model, was calculated for each cohort. This represents the degree of clustering of the raw normalised data about integer copy number values (Barnes et al. 2008). PRT data from the 1000 Genomes samples analysed is available at dbvar accession nstd159 at ncbi.nlm.nih.gov/dbvar.

### CR1 junction fragment analysis

PCR products were amplified from 5-10ng genomic DNA in a final volume of 10ul, with 0.5µl of 2.5mM of each of dATP, dCTP, dGTP and dTTP, 0.5units *Taq* DNA polymerase (KAPA Biosciences) and 0.5µl of a 10µM solution of each PCR primer. The PCR primers were 5’-AATGTGTTTTGATTTCCCAAGATCA<u>G</u>-3’ and 5’-CTCAACCTCCCAAAGGTGCT<u>A</u>-3’, with a terminal 3’ locked nucleic acid base (underlined) to increase paralogue-specificity (Latorra et al. 2003). A touch-down PCR protocol was used, with an initial denaturation step of 95°C for 2 minutes, followed by 20 cycles of 95°C for 30 seconds, 70°C for 30 seconds decreasing by 0.5°C every cycle to 60°C, and 70°C for 30 seconds. These 20 cycles were then followed by 15 cycles of 95°C for 30 seconds, 60°C for 30 seconds and 70°C for 30 seconds, then a final extension step of 70°C for 5 minutes. Products were analysed using standard ethidium bromide stained agarose gels and visualisation under ultraviolet light. Specificity of the PCR for *CR1*-B alleles was confirmed on a panel of 40 UK samples from the HRC-1 collection previously typed using PRT.

### SNP genotyping and linkage disequilibrium analysis

Genotyping for the CR1 GWAS index SNP (Lambert et al. 2013), rs6656401, and rs3818361 was carried out using KASP assays using standard protocols (LGC, Middlesex). Pairwise linkage disequilibrium was calculated using a cubic exact equation approach implemented in CubeX (Gaunt et al. 2007).

### Statistical analysis and sequence alignment

Clustal Omega was used for sequence alignment provided by the European bioinformatics Institute webserver (www.ebi.ac.uk), using default DNA options (Li et al. 2015; Sievers et al. 2011). Case-control analysis was performed using logistic regression implemented in the statistical package RStudio v.1.0 implementing R v3.2.3.

## Results

### Development and validation of PRT assays for CR1 copy number

Our first aim was to develop a simple robust approach to determine the diploid copy number of the LCRs within the *CR1* gene, which could use small amounts of DNA from large clinical cohorts. The 92% similarity between the LCRs within *CR1* and part of the *CR1L* gene allowed the design of three PRT assays to independently measure the copy number of LCR (Figure 2a).

**Figure 2.**
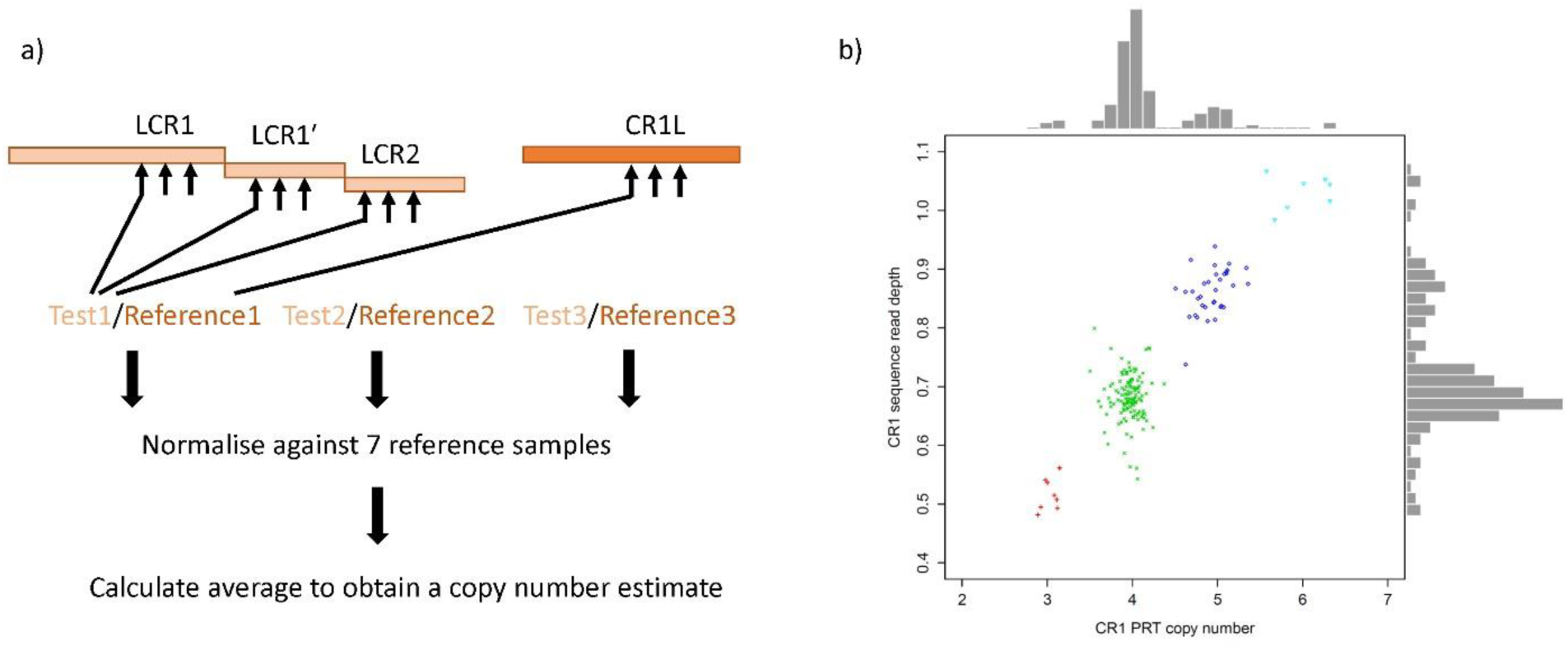
Design and validation of paralogue ratio test to detect *CR1* LCR copy number. a) Measuring copy number using the paralogue ratio test. A primer pair is designed so that it amplifies from the LCR regions (test) and also from a 92% similar region in *CR1*L (reference). Three independent tests are normalised against seven controls of known copy number, and average to obtain an estimate of copy number. b) Comparison of *CR1* PRT copy number estimate (x axis and histogram) against estimates from Illumina sequence read depth (y axis and histogram). Each point represents a different individual, points are distinguished by shape and colour indicating the final integer copy number call: red cross – 3, green x-cross – 4, blue diamond – 5, cyan triangle - 6.

Analysis of 275 samples from the HapMap collection showed high pairwise concordance (∼80%) pairwise between the three individual PRT results, allowing an average to be taken of the three PRT results for each sample as representative of the copy number of the LCR. For each sample, the copy number estimate from the PRT was compared against the copy number estimated from Illumina sequencing read depth data. The data showed a high degree of concordance, with data for both estimates clustering around integer copy number estimates. In particular, the PRT data shows clear distinct clusters, suggesting that it will call copy number accurately (Figure 2b).

Because our study was focused on the *CR1-*B duplication, distinguishing the *CR1-*A/*CR1-*B heterozygotes (diploid LCR copy number 5) from the *CR1-*A/*CR1-*A homozygotes (diploid LCR copy number 4) was particularly critical. In order to improve the reliability of distinguishing these genotypes, we developed a junction fragment PCR to specifically amplify the LCR1’ repeat. If the LCR1’ repeat was generated in the past by an equal crossover between the 98% identical LCR1 and LCR2 sequences, we would expect a switch from LCR1-like sequence to LCR2-like sequence within the LCR1’ sequence (Figure 3). Using a multiple alignment strategy on LCR1, LCR1’ and LCR2 sequences from the human reference genome GRCh37, we identified the switch point and confirmed that this is the same switch point found in early characterisations of the *CR1*-B allele (Vik and Wong 1993). Designing PCR primers flanking the switch point, with a forward PCR primer specific to the LCR2 sequence and a reverse PCR primer specific to the LCR1 will generate an amplification product from *CR1*-B alleles but not from *CR1*-A alleles (Figure 3).

**Figure 3.**
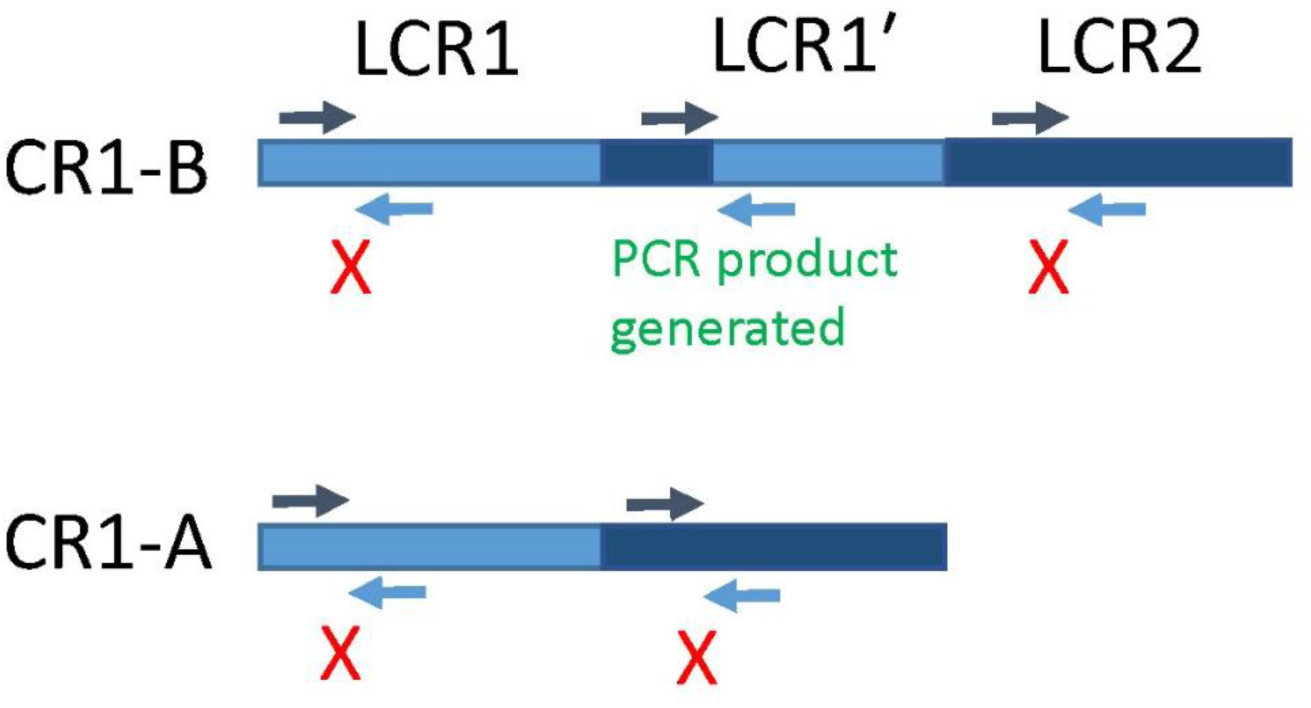
Junction fragment PCR for *CR1*-B allele. A representation of the frequent *CR1*-B and *CR1*-A alleles. Paralogue-specific primers (light-blue and dark-blue) are used to amplify specifically a breakpoint in LCR1’. Hatched repeats indicates uncertain recombinant origin.

### Association analysis of the CR1-B allele with Alzheimer’s disease

We typed 449 EOAD cases and 184 controls for copy number using our PRT approach (table 1), resulting in a dataset that showed clear clustering around integer copy numbers and good separation of clusters (Figure 4a). The Q value, a measure of clustering quality of the resulting data, was 5.56, above the minimum threshold of 4 previously suggested to be adequate for case-control studies (Barnes et al. 2008). By calling the individuals with LCR diploid copy numbers of 5 and 6 (*CR1*-B heterozygotes and *CR1*-B homozygotes respectively), we could infer *CR1*-B allele dose for each individual. We then used logistic regression with sex and ApoE4*4 genotype as covariates to test for the association of *CR1-*B allele with EOAD, assuming an additive effect of the allele. We found no evidence of association (p=0.936, table 2).

**Figure 4.**
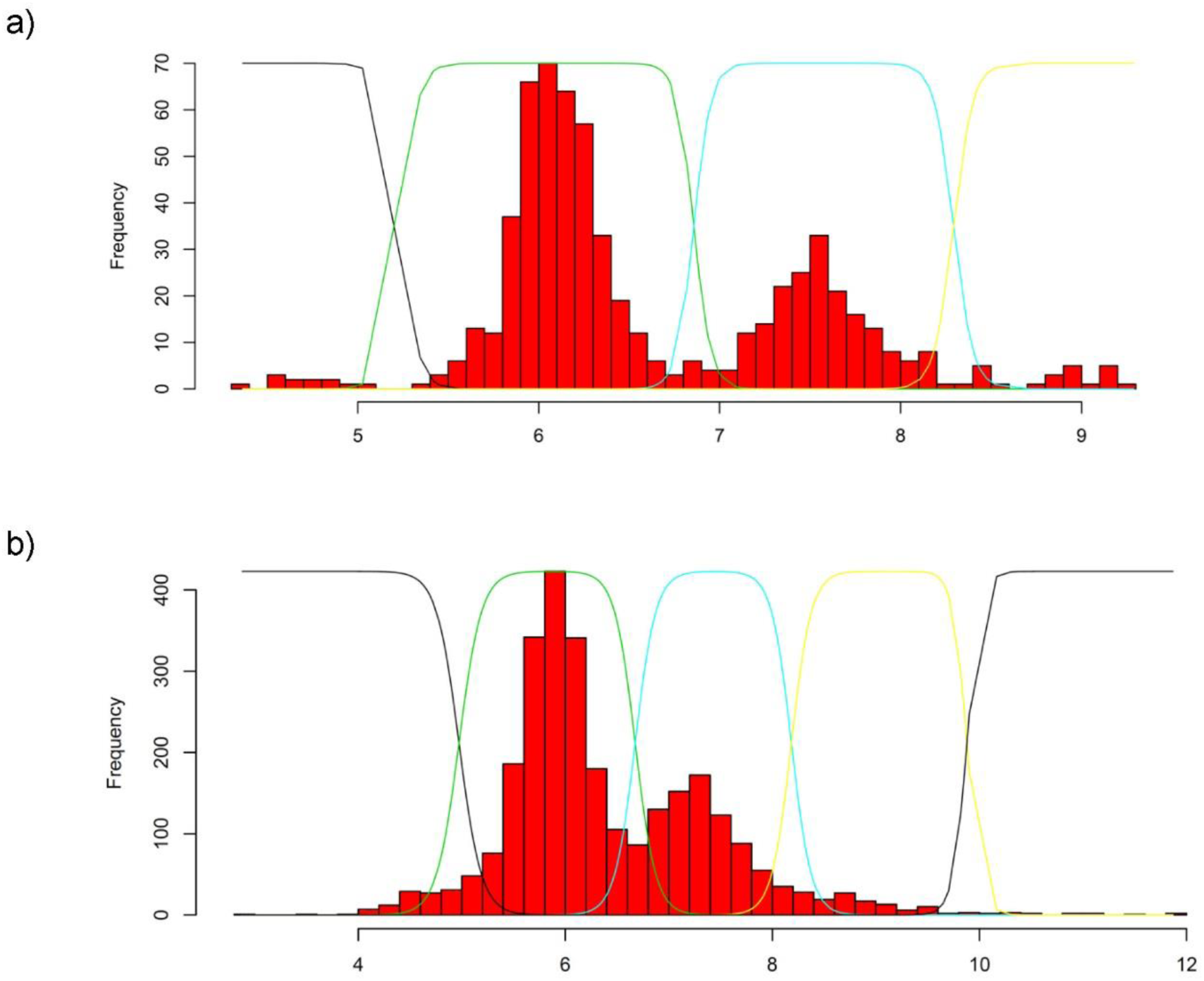
Distributions of PRT copy number data. Histograms showing the distribution of copy number values generated by PRT in a) Early onset cases and controls and b) Late onset cases and controls. X axis represents *CR1* LCR diploid copy number, with Gaussian curves superimposed to indicate the calling of integer copy number, from left to right: black – 3 copies, green 4 copies, cyan, 5 copies, yellow 6 copies and black (right hand side) >6 copies.

We then typed 1436 LOAD cases and 1175 controls for copy number using our PRT approach (table 3). The Q value for this cohort was lower (Q=4.05), and this is reflected in the clustering of values (Figure 4b), where there is significant overlap between copy number clusters. To improve calling of copy numbers 4 and 5 (i.e. *CR1*-B heterozygotes and homozygotes) the junction fragment PCR assay was also used on the LOAD cohort. We used logistic regression with sex, age and *APOE4*\*4 genotype as covariates to test for the association of *CR1-*B allele with LOAD, assuming an additive effect of the allele (table 4). We confirmed the effect of *APOE4*\*4 allele on increasing LOAD risk (p<2×10^−16^, odds ratio 3.25, 95% confidence intervals for odds ratio 2.78, 3.82), and evidence of association of the *CR1*-B allele with an increased risk of LOAD (p=0.0151, odds ratio 1.21, 95% confidence intervals for odds ratio 1.04, 1.42).

**Table 4.**
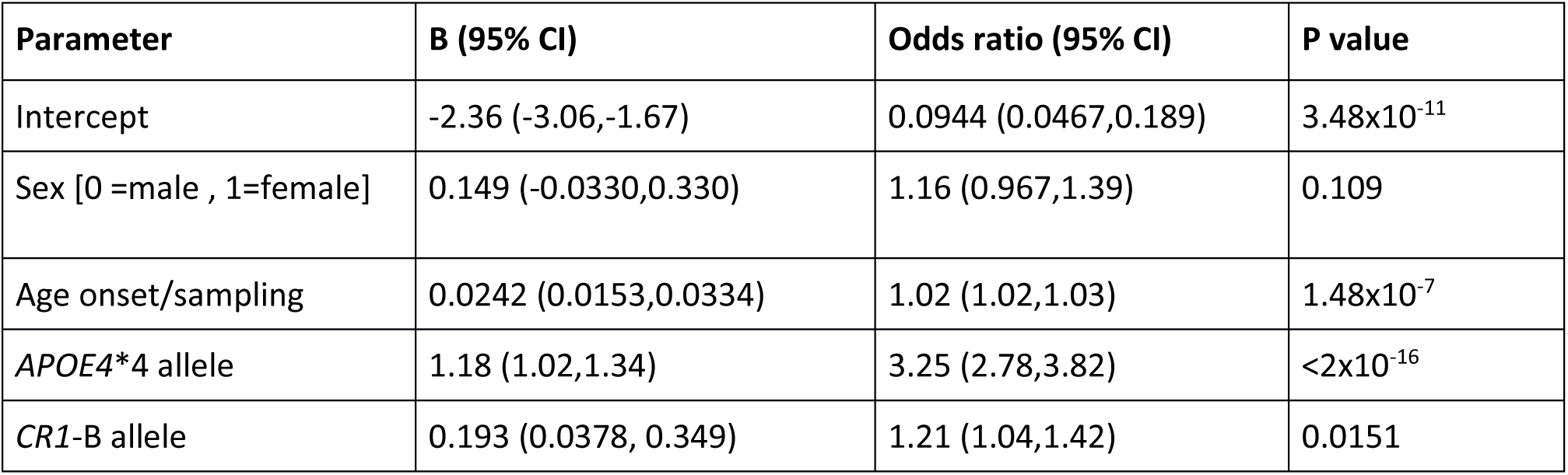
Association of *CR1*-B allele with LOAD number of cases and controls.

| **Parameter** | **B (95% CI)** | **Odds ratio (95% CI)** | **P value** |
| --- | --- | --- | --- |
| Intercept | -2.36 (-3.06,-1.67) | 0.0944 (0.0467,0.189) | $3.48 \times 10^{-11}$ |
| Sex [0 =male , 1=female] | 0.149 (-0.0330,0.330) | 1.16 (0.967,1.39) | 0.109 |
| Age onset/sampling | 0.0242 (0.0153,0.0334) | 1.02 (1.02,1.03) | $1.48 \times 10^{-7}$ |
| *APOE4* *4 allele | 1.18 (1.02,1.34) | 3.25 (2.78,3.82) | $<2 \times 10^{-16}$ |
| *CR1*-B allele | 0.193 (0.0378, 0.349) | 1.21 (1.04,1.42) | 0.0151 |

### Relationship of CR1-B with flanking SNP alleles

We next attempted to address whether the association we see explains the association at SNPs flanking the *CR1* gene seen in genomewide association studies. The strength of effect that we observe for the *CR1*-B allele (odds ratio 1.21, 1.04-1.42 95% CI) is consistent with the effect seen for rs6701713-A allele (odds ratio 1.16, 1.11-1.22 95% CI, (Naj et al. 2011)) and rs3818361-T allele (odds ratio 1.18, 1.13-1.24 95% CI, (Hollingworth et al. 2011). We analysed the pairwise linkage disequilibrium between these two SNP in a subset of the LOAD cohort (526 cases, 101 controls). We found complete linkage disequilibrium (D′=1, r^2^=1) between rs6701713 and rs3818361, indicating an A-T risk haplotype and a G-C non-risk haplotype, similar to previous findings (Mahmoudi et al. 2015). Linkage disequilibrium between both SNPs and *CR1*-B showed LD (D′=0.806, r^2^=0.576) with the *CR1*-B allele. The *CR1*-B allele occurred on the A-T risk haplotype (*CR1*B-rs6701713A-rs3818361T haplotype frequency 0.16) with *CR1*-B on the non-risk haplotype (*CR1*B-rs6701713G-rs3818361C) infrequent at a frequency of 0.03. This pattern of LD is consistent with the *CR1*-B allele explaining the association observed at the flanking SNP alleles.

If the association of LOAD with the *CR1-*B allele explains the association at flanking SNP alleles then we would expect a stronger association of disease with the *CR1* duplication compared to the SNP alleles. We analysed a subset of our cohort that had genotypes for rs6701713 and rs3818361 as well as *CR1*B genotype, using the same model used to analyse the full LOAD cohort. Due to limited DNA availability, this subset was small (478 cases and 96 controls) and underpowered to detect evidence for an association *a priori*. Nevertheless, the association with *CR1*-B was stronger (p=0.067, OR 1.52, 95%CI 0.99-2.44) than with either rs6701713A or rs3818361T (p=0.56, OR=1.13 95%CI 0.76-1.72), and incorporating *CR1*-B rather than rs6701713A or rs3818361T into the logistic regression model provides a better fit to the data (Akaike’s information criterion=469.99 vs 473.26).

## Discussion

In this study we develop a new approach to measuring the copy number variation of LCRs within the *CR1* gene using the paralogue ratio test, together with a junction fragment PCR specific for the *CR1*-B allele that carries three copies of the LCR. We typed a case-control cohort of early-onset Alzheimer’s disease and a case-control cohort of late-onset Alzheimer’s disease for copy number of the LCR within the *CR1* gene, inferring the *CR1*-B genotype. An association study of the *CR1*B allele with EOAD showed no evidence of association, but an association study of the *CR1*B allele with LOAD provided evidence of association of the *CR1*-B allele with increased LOAD risk, in agreement with previous studies. We also show that the *CR1*-B allele is on the LOAD risk haplotype identified by SNP-based GWAS, and that the *CR1*-B allele shows stronger evidence of association with LOAD than the risk SNP alleles identified by GWAS.

One limitation of the PRT approach is that it reports diploid copy number – i.e., the copy number summed over both alleles, rather than the true genotype. So, for example, a copy number of 5 could be a 3 allele and a 2 allele, or a 4-1 or a 5-0. We assumed that all 5 copy individuals were 3-2, that is *CR1*-B/*CR1*-A and all 6 copy individuals were 3-3 *CR1*-B/*CR1*-B not 4-2 *CR1*-A/*CR1*-D. This could be an incorrect assumption leading to an overestimation of the frequency of the *CR1*-B allele. Previous work has shown, by western blotting of the CR1 protein, that in three out of eight individuals the true genotype was *CR1*-A/*CR1*-D not *CR1*-B/*CR1*-B, suggesting that a significant proportion of 6 copy individuals may not be homozygous for *CR1*-B as we assume (Brouwers et al. 2012). In order to assess the validity of our assumption in our population, we estimated the population allele frequencies from the diploid copy number data of the EOAD cohort using the R script CNVice, which simultaneously tests for any departure of the inferred genotype frequencies from Hardy-Weinberg equilibrium (Zuccherato et al. 2017). The relative probability of a genotype given the individual’s copy number was also calculated from these results. We found that 4-copy individuals had a 99.2% probability of being *CR1*-A homozygotes, 5-copy individuals had a 100% probability of being *CR1*-A/*CR1*-B heterozygotes and 6-copy individuals had a 100% probability of being *CR1*-B homozygotes, supporting our assumptions used in this study.

The mechanistic basis for the association of the *CR1*-B allele with LOAD remains unclear. It has been suggested that, because of its extra C3b-binding site, the CR1-B protein is more effective at inhibiting C3b complement fragments, leading to a reduction in C3b-mediated opsonisation of Ab1-42 fragments (Brouwers et al. 2012). However, in brain the CR1-B isoform is expressed at lower levels than *CR1*-A and is probably associated with increased complement activation; indeed, complement system is activated by Ab (Hazrati et al. 2012; Mahmoudi et al. 2015; Rogers et al. 1992). Future studies need to focus on the functional effect of *CR1*-B allele *in vivo*, in combination with structural studies to determine the difference in structures between the different *CR1* alleles.

## Acknowledgements

We would like to thank Daniel Zadik for help and advice, and Rachael Madison for technical support. All authors have read and approved the final manuscript. The ARUK Consortium was funded by Alzheimer’s Research UK, with the following members:

## Tables

**Supplementary Table 1.**
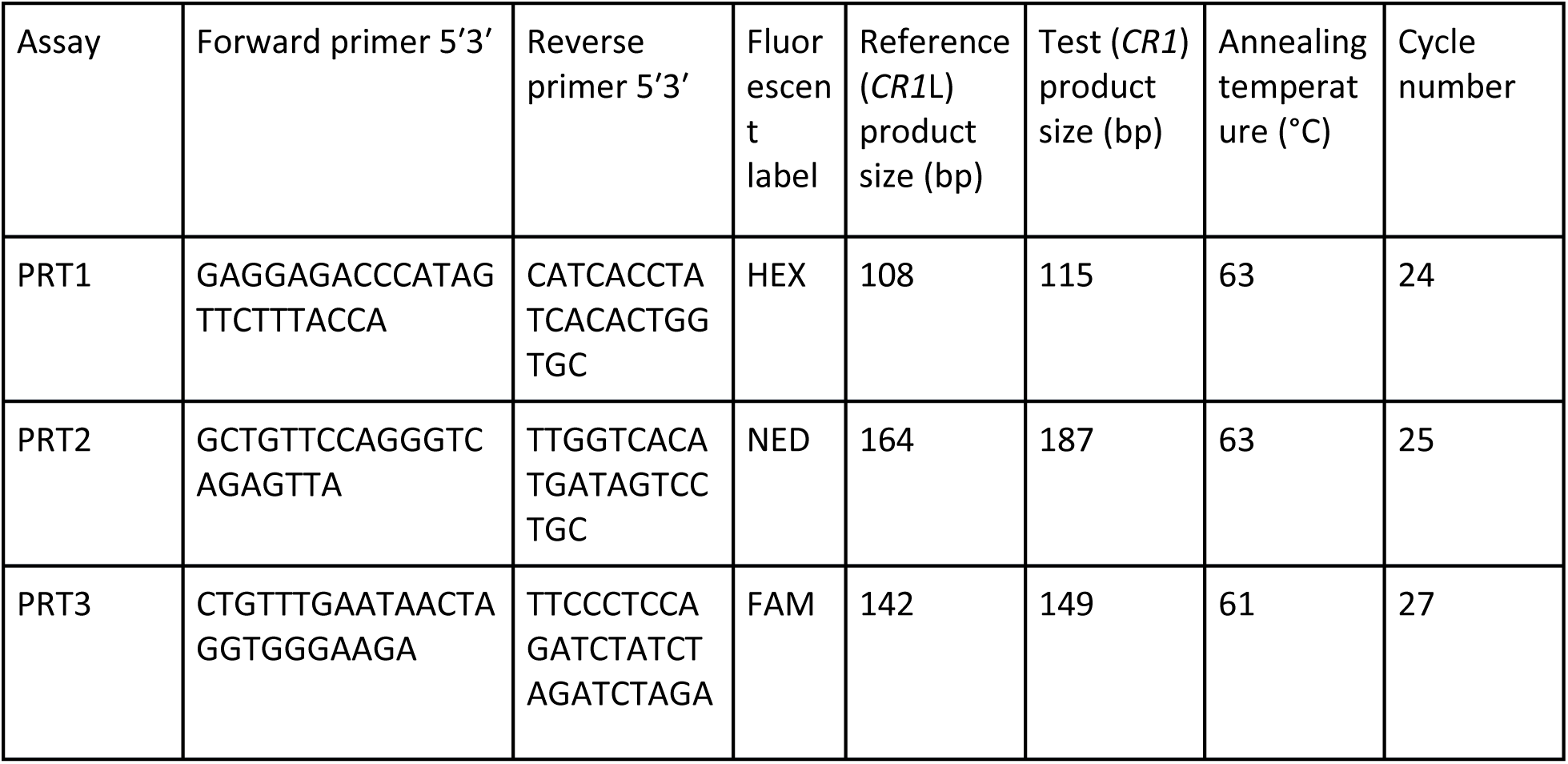
PRT assays for *CR1* LCR copy number.

**Supplementary table 2.**
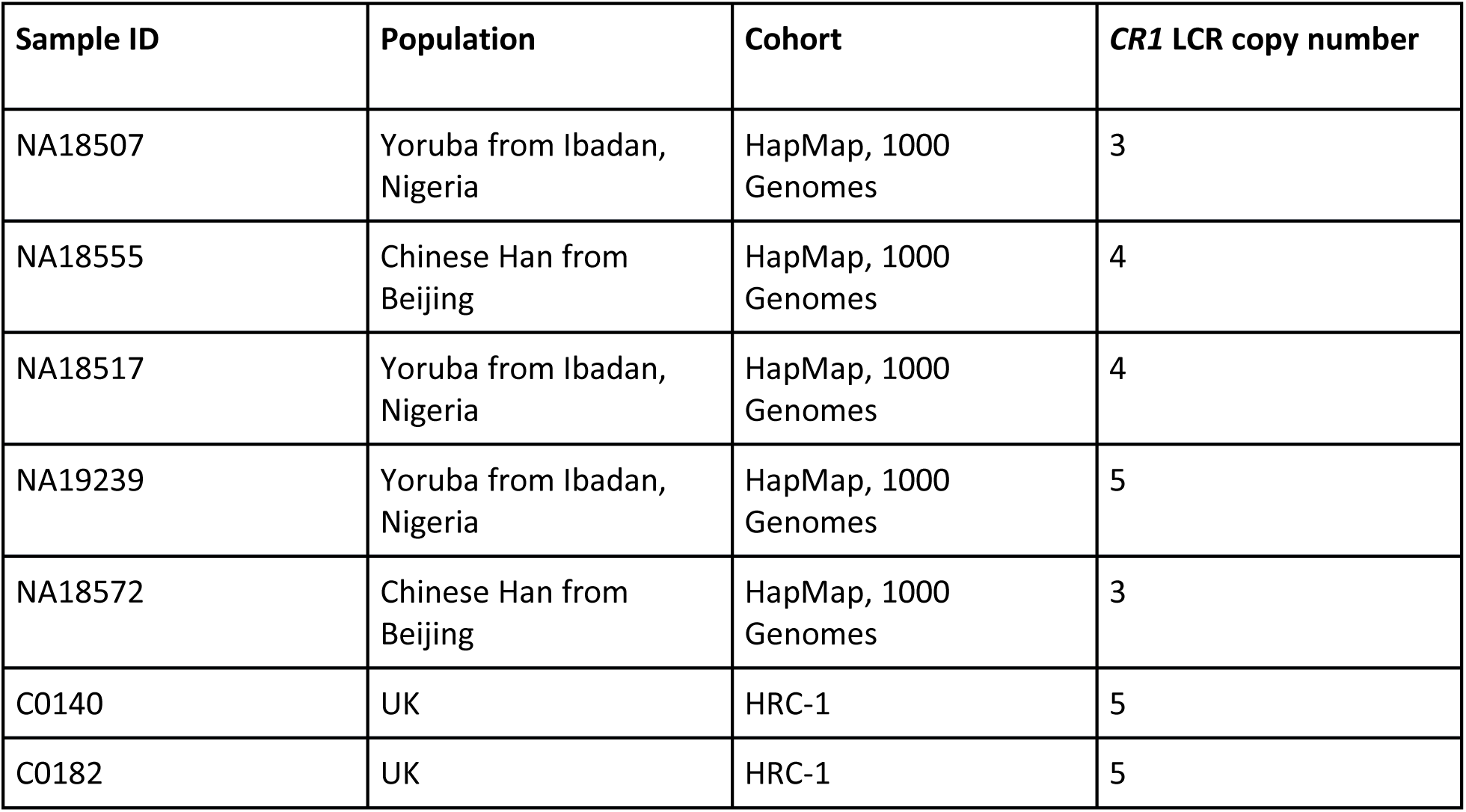
Control samples for *CR1* paralogue ratio test.

